# Cloud Bursting Galaxy: Federated Identity and Access Management

**DOI:** 10.1101/506238

**Authors:** Vahid Jalili, Enis Afgan, James Taylor, Jeremy Goecks

## Abstract

**Motivation:** Large biomedical datasets, such as those from genomics and imaging, are increasingly being stored on commercial and institutional cloud computing platforms. This is because cloud-scale computing resources, from robust backup to high-speed data transfer to scalable compute and storage, are needed to make these large datasets usable. However, one challenge for large-scale biomedical data on the cloud is providing secure access, especially when datasets are distributed across platforms. While there are open Web protocols for secure authentication and authorization, these protocols are not in wide use in bioinformatics and are difficult to use for even technologically sophisticated users.

**Results:** We have developed a generic and extensible approach for securely accessing biomedical datasets distributed across cloud computing platforms. Our approach combines OpenID Connect and OAuth2, best-practice Web protocols for authentication and authorization, together with Galaxy (https://galaxyproject.org), a web-based computational workbench used by thousands of scientists across the world. With our enhanced version of Galaxy, users can access and analyze data distributed across multiple cloud computing providers without any special knowledge of access/authorization protocols. Our approach does not require users to share permanent credentials (e.g., username, password, API key), instead relying on automatically-generated temporary tokens that refresh as needed. Our approach is generalizable to most identity providers and cloud computing platforms. To the best of our knowledge, Galaxy is the only computational workbench where users can access biomedical datasets across multiple cloud computing platforms using best-practice Web security approaches and thereby minimize risks of unauthorized data access and credential use.

**Availability and Implementation:** Freely available for academic and commercial use under the open-source Academic Free License (https://opensource.org/licenses/AFL-3.0) from the following Github repositories: https://github.com/galaxyproject/galaxy and https://github.com/galaxyproject/cloudauthz

## I. Introduction

Genomics has become an essential tool in many biomedical areas, including developmental biology, human evolution, and precision medicine. As DNA sequencers become more affordable and pervasive, increasing volumes of sequencing data are being generated and stored on a variety of computing platforms. Genomics is expected to be an exabase-scale big data domain by 2025, posing data acquisition and storage challenges on par with astronomy, YouTube, and Twitter—the other major generators of big data [1]. Large and distributed genomics data poses key challenges for making effective use of the data, especially for the increasingly important tasks of data integration and joint analysis.

The emergence of cloud-based computing platforms such as Amazon Web Services (AWS) and Microsoft Azure have paved the path for online, scalable, cost-effective, secure, and shareable big data persistence and analysis with a growing number of researchers and laboratories hosting (publicly and privately) their genomics big data on cloud-based services [2]. Most computational analyses of genomics data requires complex workflows that include many steps and analysis tools. Computational workbenches such as Galaxy [3] and GenomeSpace [4] make it simple to create and execute analysis workflows. Consequently, these systems need to seamlessly access cloud-based genomics data to execute workflows.

Perhaps the first challenge of integrated genomic data analysis on cloud-based platforms is getting access to the data. Data access involves authentication—verifying a user’s identity—and authorization—determining what data/resources a authenticated user can access. Given that genomics data, especially from clinical samples, is often highly sensitive—as it may contain protected health information (PHI) or personally identifiable information (PII) [5]—protocols are needed to implement secure authentication and authorization.

Several systems such as GenomeSpace and Globus [6] use cloud-based platforms for data storage and transfer. However, they rely on storing permanent user credentials for a given cloud platform. For instance, to transfer data to/from a cloud platform, Globus prompts for user credentials with the provider which are then cached and used to access the resources. Delegating a user’s private credentials to a third-party service is a privacy and security risk for the user and a liability risk for the service. Additionally, revoking these access credentials requires manual intervention, and obtaining credentials requires users to be familiar with the technical details of the given cloud provider, hindering adoption by users with limited technical knowledge. These challenges highlight the need for a robust solution to securely delegate access to cloud-based genomics data.

We have developed an approach for federating identity and access management and implemented it in the Galaxy framework. Our approach enables a secure and seamless delegation of privileges without sharing users’ login and/or access credentials. With this approach, Galaxy users can securely access genomics data across a wide variety of cloud computing platforms. The advantages of our approach are twofold: (a) leveraging OpenID Connect (OIDC) protocol to allow users to login to Galaxy using existing third-party identities, and ability to securely grant Galaxy authorization to access genomic data on the cloud. Leveraging the OIDC protocol for user authentication paves the path for the Galaxy-as-service model [7] where a user can login to all Galaxy servers using a single identity. In addition to simplifying the login process, this model protects users’ identity and credentials should any Galaxy server suffer a security breach.

The second advantage of this approach is that a user can securely grant Galaxy authorization to access genomic data on the cloud. Previously, users needed to share their permanent cloud credentials with Galaxy [8], which is problematic because those credentials grant Galaxy the same privileges as the user, Galaxy must store and secure those credentials, and users must manually obtain those credentials. We have developed a new approach that leverages on-demand and automatic generation of temporary access credentials to assume minimum delegated privileges. This approach reflects best-practice approaches on the web, and it minimizes risk and difficulty for users when accessing data on the cloud. This approach was implemented for multiple cloud providers in a reusable library called *CloudAuthz* (https://github.com/ galaxyproject/cloudauthz).

The described approach is integrated into the Galaxy application framework (https://galaxyproject.org), making it possible for Galaxy users to securely access and analyze cloud-based genomics data. Galaxy users can access cloud data that they own or have access to by specifying a provider name (e.g., AWS) and a resource name (e.g., an Amazon Simple Storage Service (S3) bucket), but share neither login nor access credentials (see section II). Being able to perform these steps without requiring the user to supply their permanent access credentials has the following advantages:

- Galaxy does not ask for or store user access credentials, and users can authenticate using available Web identity providers such as Google. Thus our approach is both user-friendly and secure;
- User identity is authenticated via security tokens that cannot be exploited to impersonate them.
- Authorization to data follows the principle of least privilege, so Galaxy can read/open given datasets but does not have access to other user data or compute resources;
- Delegated privileges for Galaxy, which are defined by short-term authentication/authorization tokens, are independent from a user’s credentials, hence their scope can be restricted independently;
- Can leverage the role-based access control model [9], which enables segregating duties and provide Galaxy with the least privileges required for data access;
- The short-term authentication/authorization tokens issued for Galaxy are refreshed automatically and can be revoked by the user, either from the identity provider (e.g. Google)or the cloud platform;
- Privileges for a Galaxy server cannot be used by a different client (e.g., another Galaxy server, or a different web app). This is guaranteed by OIDC protocol.

### A. Motivating Application

Cloud-based services have become a ubiquitous storage solution due to their scalability, availability, and cost efficiency, which make them an ideal storage solution for the genomics big data that are commonly studied in a collaborative setting. Accordingly, a growing number of datasets are publicly hosted on cloud service providers. For instance, *Tabula Muris* is a single-cell transcriptomic dataset comprising more than 100,000 cells of 20 organs and tissues of *Mus musculus* [10] and it is publicly hosted on AWS (accessible through the following Amazon Resource Name: arn:aws:s3:::czb-tabula-muris). In addition to *Tabula Muris*, 87 additional datasets exist from various disciplines that are all publicly available via the AWS *registry of open data* (https://registry.opendata.aws). Among them Éare *The Human Microbiome Project* (arn:aws:s3:::human-microbiome-project), *The International Cancer Genome Consortium* (arn:aws:s3:::oicr.icgc.meta/metadata), *Nanopore Reference Human Genome* (arn:aws:s3:::nanopore-human-wgs), and *1000 Genomes* (arn:aws:s3:::1000genomes).

A common scenario that demonstrates the need for robust authentication and authorization to cloud-based genomics data is joint analysis of public and private datasets. For instance, an active area of research in precision oncology is using omics data to statistically learn biomarker signatures to guide selection of therapies [11], [12], [13], [14] most likely to be effective for a particular tumor. In these studies, both private as well as public datasets such as The Cancer Genome Atlas (TCGA) require authorized access. Currently this kind of research requires copying private and public datasets on the same institutional computing cluster or cloud computing platform and running analyses on that cluster/platform. Moving omics data is costly, difficult, and can be insecure depending on how access credentials are used. Our approach greatly simplifies joint analyses by providing a secure way for users to access data on one or more clouds. With the described new feature of Galaxy, users can securely access and combine both private and public datasets on a single Galaxy server where they can be analyzed together. This server can live behind an institutional firewall or on a commercial cloud computing platform and hence provide flexibility about where the final analysis is run.

Large-scale collaboration is another scenario where secure cloud-based authorization to genomics data is critical. For instance, collaborating labs across different institutes can host their data on the cloud and grant each member of those labs read (and write) access to the data. The challenge, however, is the ability to readily access that data for analysis. Typically, this is accomplished by either downloading the data onto on-premises resources or “mounting” it on cloud-based compute platforms. With Galaxy being widely adopted as a scalable, transparent, and reproducible data analysis platform, it is essential to enable Galaxy users to load their cloud-hosted, private data into their Galaxy *history* in an attestable and auditable manner. Accordingly, a resource owner can share cloud-hosted data with a user who is authenticated using their social or institutional identities. The user can then login to Galaxy using their specified identity, and request copying shared data into their *history*. Having analyzed the data, the user can request copying analysis results from the Galaxy *history* to the cloud-hosted storage, which enables them to share the analysis results with their collaborators. An illustration of this scenario is given in Figure 1, and a detailed discussion of the method is available in Section II. For example, it is now possible to authenticate with Galaxy using a Google identity. Given a one-time, out-of-band setup where that identity is associated with an AWS S3 bucket role, the authenticated Galaxy user can seamlessly download and upload data to a private S3 bucket. All this is done without ever prompting the user for their AWS credentials.

**Fig. 1.**
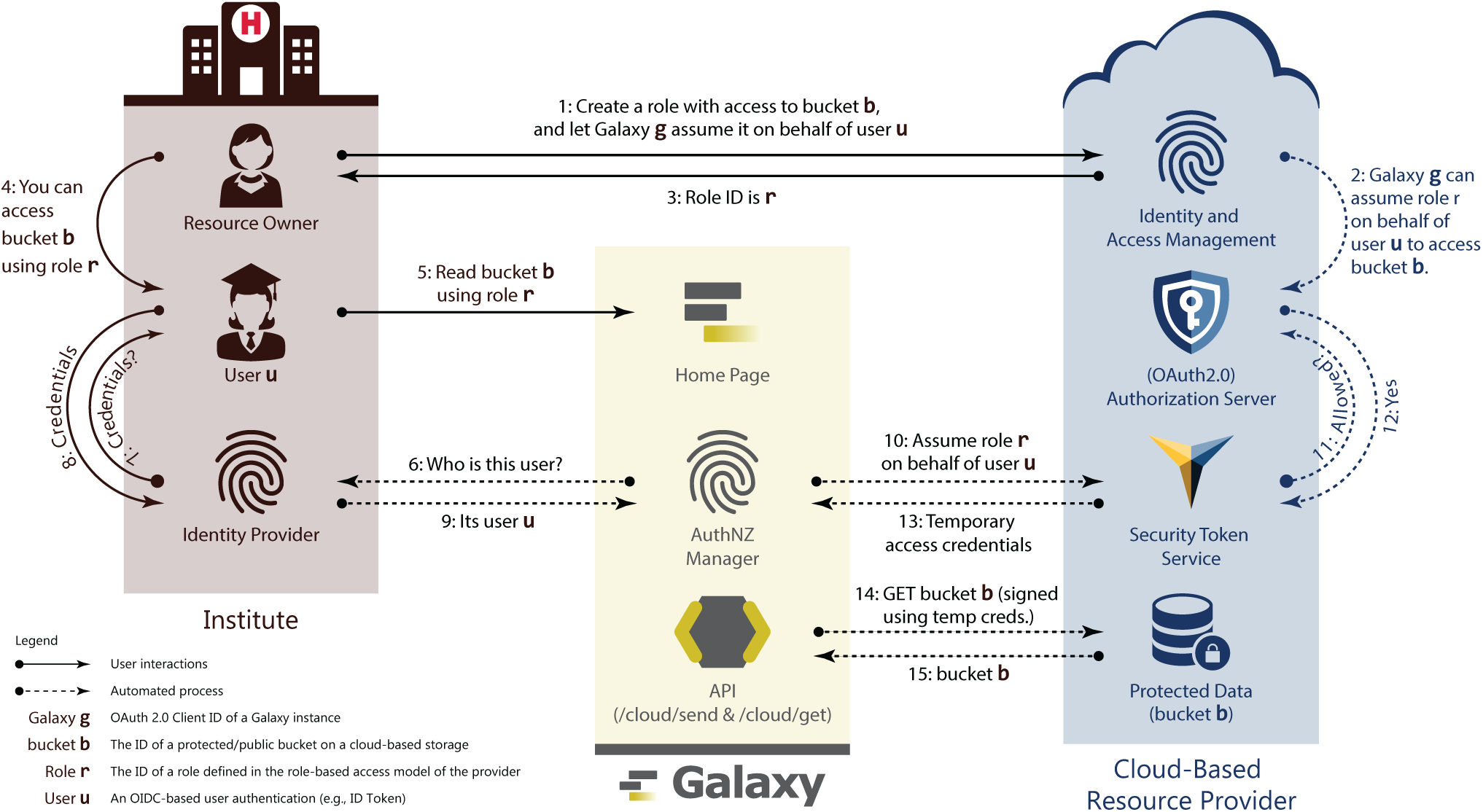
Galaxy adopts and integrates best-practice Web protocols to access secured data stored on cloud platforms (discussed in details in Section II). In this approach, a *resource owner* shares protected data with collaborators (*User*) leveraging the role-based access model [9] and OpenID Connect protocol (OIDC). Accordingly, a resource owner defines a *role* with (read or write) access to protected data (e.g., see Figure 3), and specifies a Galaxy instance (defined using OIDC *audience* ID) that can assume the *role* upon presenting the user’s identity token issued by their specified institute (OIDC IdP) for that Galaxy instance (e.g., see Figure 4). Upon successfully assuming the *role*, Galaxy receives cloud-provider-specific temporary credentials, and uses them to sign API requests to protected data. Note that following the OIDC requirements, all the discussed communications are TLS-protected (see Section III-A). Additionally, a *resource owner* and *user* are not required to belong to a same trust group (e.g., institute).

## II. Methods

Linking a cloud-based storage to Galaxy without requiring user credentials is realized by leveraging the OIDC protocol and the *CloudAuthz* library; this is implemented as a two-step authentication and authorization procedure. Authentication allows a user’s identity to be validated while authorization verifies the privileges the given user has. Galaxy currently supports user authentication through ¡3000 identity providers, and supports secure authorization delegation for AWS, Microsoft Azure, and Google Cloud Platform (see Figure 2).

**Fig. 2.**
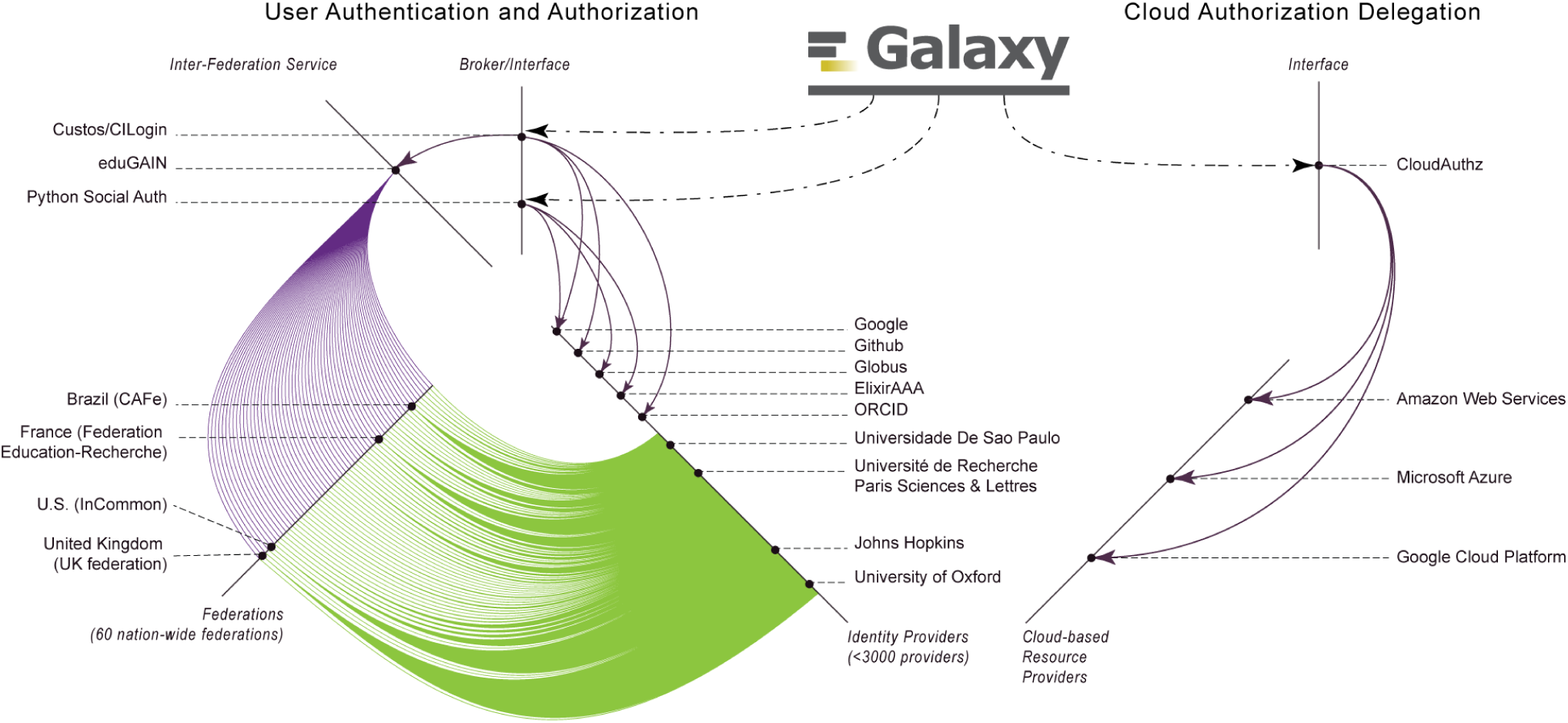
Galaxy has enabled users to login using their identities with a wide-range of identity providers, spanning from Google, Github, ORCID, ElixirAAA, and Globus, to ¡3000 world-wide educational institutes. Accordingly, Galaxy leverages CILogon and Python Social Auth for users authentication, and these brokers interface with a number of (social and institutional) identity providers, and CILogin interfaces with *eduGAIN* that federates 60 nation-wide federations of educational identity providers. For instance, top-4 federations in terms of the IdPs they integrate are *United Kingdom (UK federation), U.S. (InCommon), France (Fédération Éducation-Recherche)*, and *Brazil (CAFe)*, and one institute per federation is highlighted in the figure. Additionally, Galaxy leverages CloudAuthz to obtain authorization to cloud-based resource providers, such as AWS, Azure, and Google Cloud Platform.

### A. User authentication

Galaxy leverages the *authorization code flow* of the OIDC protocol to authenticate (and authorize) a user. In this flow, a user’s identity is first verified by an IdP, then Galaxy receives security tokens (e.g., ID token and access token) from the IdP, which contains claims about the authentication and authorization of the user. The tokens are represented in cryptographically-signed JavaScript Object Notation (JSON) Web Tokens, JWTs, which ensures their integrity and immutability. In general, this flow is a two-step procedure described as follows.

First, the admin of a Galaxy instance sets up the instance for the *authorization code flow* by registering the instance with the IdP, and obtains security credentials for the instance (e.g., *Client ID* and *Client secret* as provided by Google). These credentials are used to ensure the authenticity of communications between the parties. For instance, an identity token issued for a user contains an *audience* claim (the *Client ID* of that Galaxy instance), which ensures that the token is issued for and can be used by the specified audience only.

Second, Galaxy authenticates a user (who wants to login to Galaxy using their third-party identity) by sending a request to an IdP. Among other information, the request incorporates:

- Security tokens of the Galaxy instance obtained when registering the instance (e.g., client ID and client secret as used with Google);
- Redirect URL; to be called upon successful authentication;
- Anti-forgery claims (e.g., *state* and *nonce* to prevent respectively cross-site request forgery (XSRF) and replay attacks).

Upon a successful authentication, an IdP sends an authorization *code* and the *state* token to the Galaxy instance. The Galaxy instance uses the state token to validate the authenticity of the redirect message, and associate the authorization code with a user of the Galaxy instance. Then the Galaxy instance exchanges the authorization code for an ID token and a refresh token (which can be used to refresh an expired ID token) from the IdP.

### B. Authorization Grant

In general, cloud-based resource providers leverage the role-based access control (RBAC) model [9] to grant authorization. However, each resource provider implements a proprietary procedure to authorize a client to assume a *role*. For instance, while an AWS role can be assumed using access key and secret key, a client has to provide subscription ID, client ID, client secret, and tenant ID to assume a role (service principal) on Microsoft Azure. However, the presented method is generic and can be used on any RBAC and OIDC-compliant resource provider. The following sections explain the method on AWS and Azure for defining and assuming a role, where a role is defined via the resource provider’s web portal and it is assumed in Galaxy leveraging *CloudAuthz*.

#### 1) AWS Temporary Credentials

An AWS *role* is an identity that can be assumed by an OIDC Relying Party (RP)—Galaxy in our scenario—on behalf of a user who is authenticated by an IdP. A role has certain permissions to specific resources (e.g., read access to a S3 object) that are defined using policies attached to it (e.g., see Figure 3). Through the AWS identity and access management web portal, a resource owner defines a role and a policy, and attaches the policy to the role. The resource owner then defines a *trust* relation for the role, which defines the principals who are authorized to assume the role. The trust relation is defined using the *audience* ID of an RP, and authorizes the RP to assume the role on behalf of an IdP-authenticated user (e.g., see Figure 4). The *audience* ID is a required claim of an ID token that AWS security token service (STS) uses to assert if the token presented for assuming a role is issued for the RP defined in the trust relation. This mechanism prevents assuming a role using an ID token that is issued for a RP other than the one defined in the role’s trust relation.

**Fig. 3.**
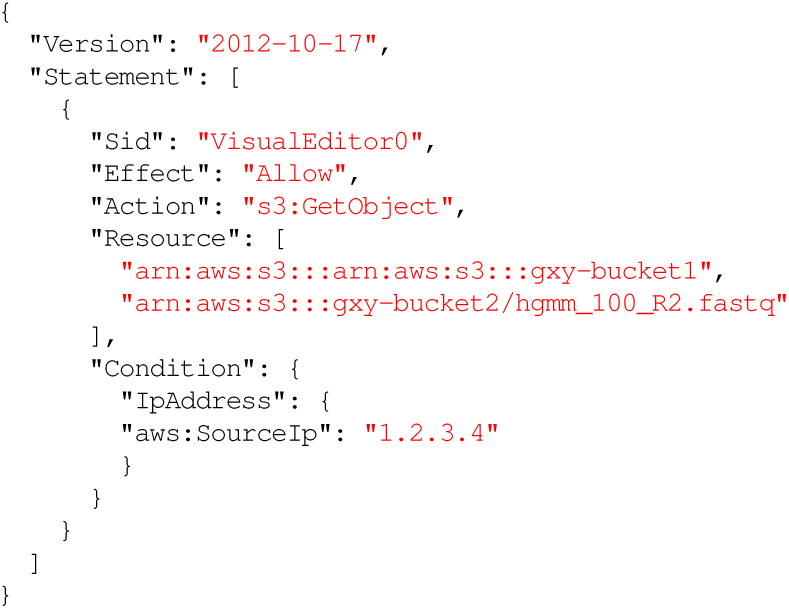
A sample of an AWS policy, which can be attached to a role to enable it to *retrieve* (`“Action”: “s3:GetObject”`) all the objects in the bucket `gxy-bucket1` and only the object `hgmm_100_R2.fastq` from bucket `gxy-bucket2`, if the request is made from a server with `1.2.3.4` IP address. *Sid*: statement ID.

**Fig. 4.**
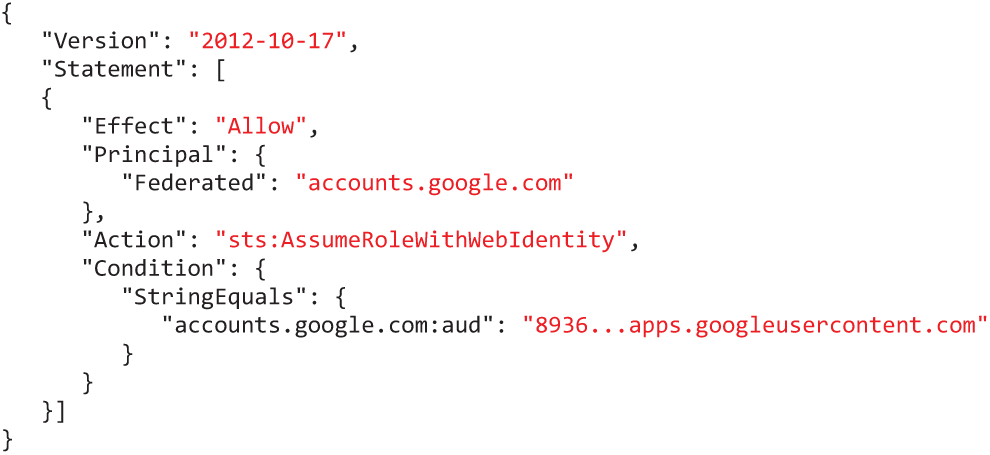
An example of a trust relationship defined for an AWS role, which allows a Galaxy instance, identified by the `8936`…apps.googleusercontent.com (part of the client ID), to assume the role in exchange of a user’s ID token issued for that Galaxy instance by Google.

A user defines an AWS role for Galaxy using its Amazon resource name (ARN). Galaxy assumes the role by submitting a request to Amazon STS, which contains the role ARN and the user’s ID token. AWS STS asserts the authenticity of the request by verifying with the IdP if the ID token is not expired and is issued for the audience specified in the ID token, and if the audience is trusted to assume the role. After a successful validation AWS STS responds to Galaxy, which among other information includes *access key ID, secret access key*, and *session token*. These credentials can be used to assume delegated privileges (e.g., read an AWS bucket) as defined in the policy attached to the role (see Figure 5). The temporary credentials are automatically refreshed by Galaxy, and can be restricted (update policy) and revoked by the resource owner.

**Fig. 5.**
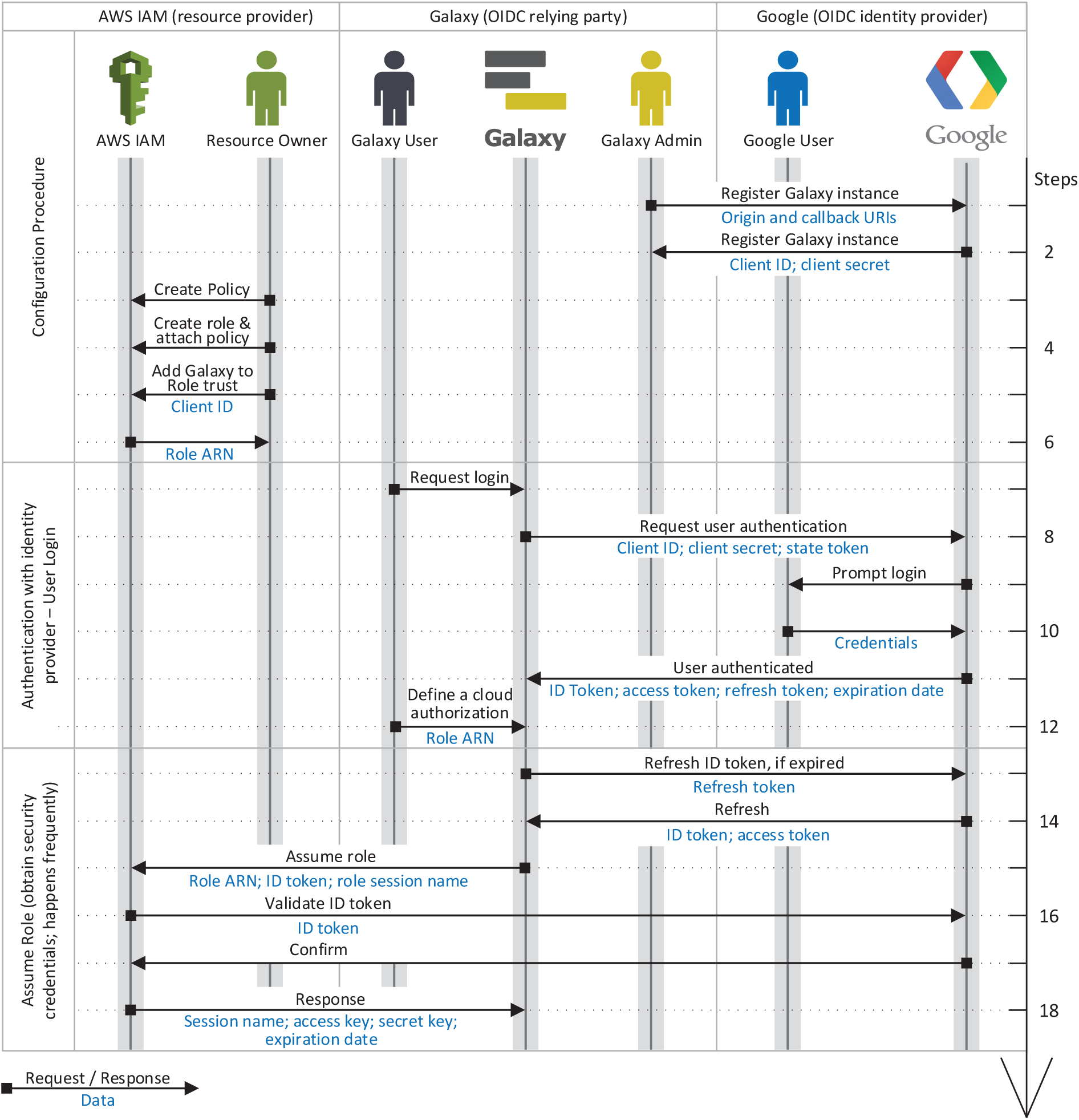
Identity federation and authorization grant flow in the proposed method for AWS. The flow is a three-step procedure; first, a Galaxy admin registers the instance as an OIDC *client* with and OIDC IdP (e.g., Google), and a resource owner configures an AWS *role* that can be assumed by the Galaxy instance (specified using its OIDC *client ID*, see Figure 4) to perform a certain operations on their resources (see Figure 3). Second, the user logs in to Galaxy (either as a new user, or in association with their existing account) using their identity with the OIDC IdP with which the Galaxy instance is registered as a *client* (e.g., Google), and they define a cloud authorization record using the AWS role ARN the resource owner has shared with them. Third, Galaxy communicates with Amazon Secure Token Service (STS), presents all the necessary information to assume the role, and obtains *access key, secret key*, and *session token*, which can be used to sign API requests to AWS resources. Note that a *Galaxy user* and a *Google user* can refer/belong to a same person; however, they are not necessarily the same *identities*, as a *Galaxy user* can be associated with multiple identities on different IdPs.

#### 2) Azure Service Principal

Azure resource manager lever-ages RBAC model [9] to enforce permissions. The operations an Azure *role* is authorized to perform are defined by its *permissions* and *scope*. Azure defines several built-in roles (e.g., the *Storage Blob Data Reader* role has read access to data and containers of Blob storage), and allows defining custom roles using Azure PowerShell or Azure CLI. An Azure role is assigned to a *security principal*, which defines a user, a group of users, or a *service principal*. A service principal is an identity used by applications or services. Accordingly, to authorize a Galaxy instance to access protected Azure resources, the resource owner defines a service principal and assigns an appropriate role to it.

A client can assume a service principal leveraging the *client credentials grant* flow of OAuth 2.0 protocol. This is a non-interactive flow and it is specifically designed for application-to-application communication. In this flow, client authentication (*ID* and *secret* of the service principal) is used as the authorization grant. Accordingly, neither the resource owner nor a Galaxy user is asked for a consent when client attempts to assume a service principal by a client, hence, this flow should be established between confidential clients only.

To assume an Azure role, a client requests an access token from Azure’s authorization server using its client credentials (see Figure 6). Upon a successful client authentication, the authorization server issues an access token for the client. The access token is issued for the application, independent from a user, and grants the client with privileges as defined by the role attached to the service principal. Following the client credentials grant flow, Azure’s authorization server does not provide a *refresh token*; hence, an expired access token is refreshed by repeating the authorization process. Additionally, the authorization is revocable by removing the service principal, or changing its *secret*, or updating the role’s assigned to it.

**Fig. 6.**
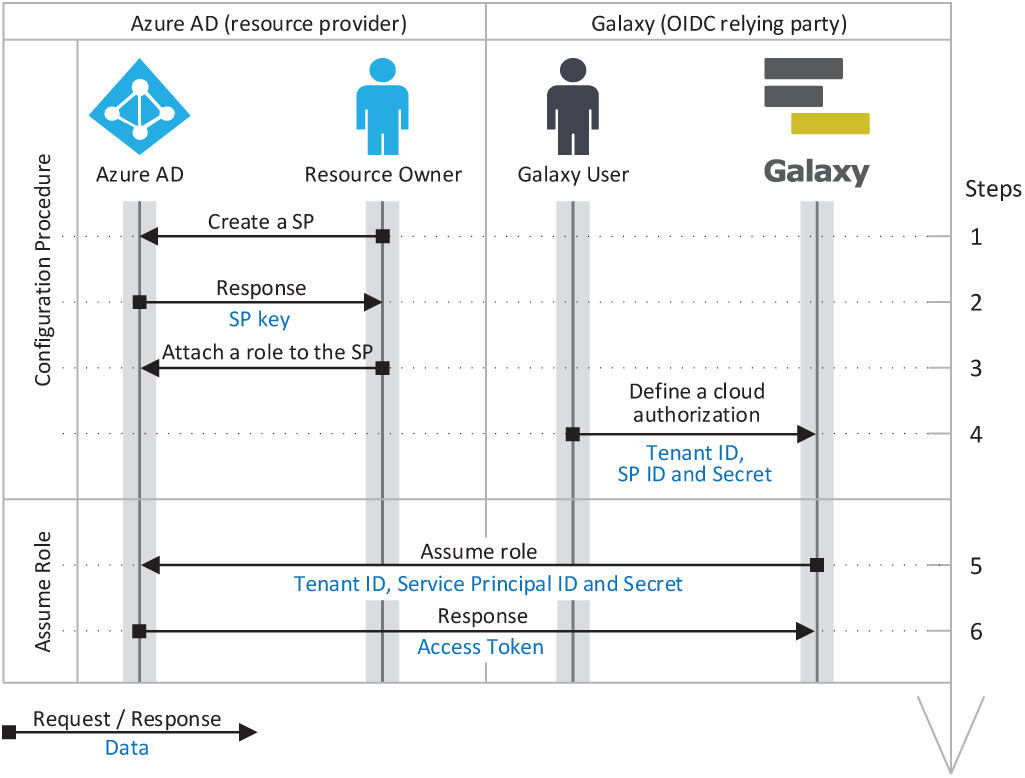
Identity federation and authorization grant flow for Azure. Accordingly, a resource owner defines a *service principle* (SP) and assigns a *role* with necessary permissions (e.g., read a bucket) to it, and obtains its *client ID* and *client secret*, and shares them, along with *tenant ID*, with a Galaxy user. (A resource owner and a Galaxy user can potentially be the same person; however, they are not necessarily the same *identities*.) The user can then define a cloud authorization record in Galaxy using the *tenant ID, client ID*, and *client secret*, which Galaxy can use to assume the role and obtain OAuth2.0 access token. Note that, this is the *client credentials grant* flow of OAuth 2.0 protocol that allows assuming a role using the aforementioned information only, and without needing for user’s OIDC identity token (unlike the flow presented for AWS in Figure 5).

## III. Results

Galaxy federates users identity and authentication using the OIDC protocol, which is the current industry standard. With this model, an identity provider authenticates a user to a Galaxy instance using temporary identity token. Galaxy then uses this token to obtain cloud-native credentials and use resource provider’s API to access protected resources (a detailed discussion is postponed to Section III-B2). To continue operating on user’s behalf beyond the validity of the initial token, Galaxy automatically refreshes the token as a trusted party. To simplify the process, manual intervention for users is minimized (see figures 1, 5 and 8) and the tokens/secrets are never handed-out to end users.

**Fig. 7.**
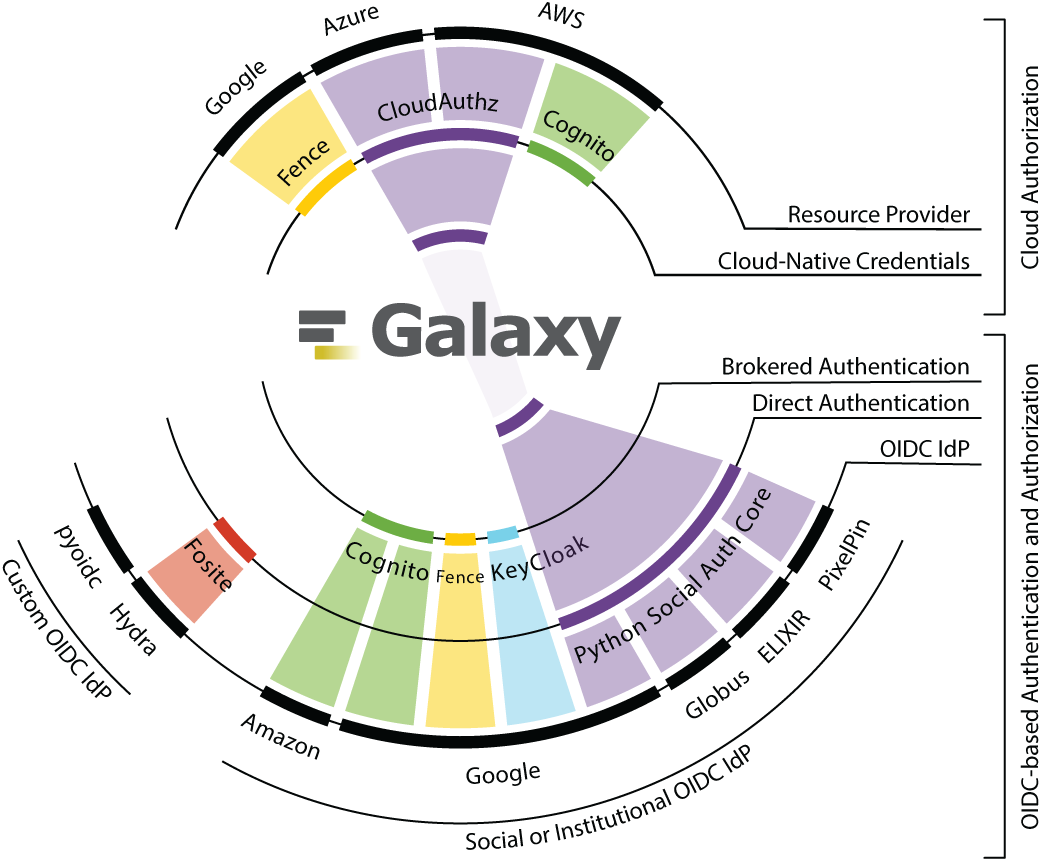
Illustrates a subset of available methods and implementations for user authentication and authorization grant to cloud-based resource providers, and the back-ends each method supports. The figure is scoped to only OIDC-based authentication and authorization grant using cloud-native credentials. The method and implementations we use in Galaxy are highlighted in *purple*, which are Python Social Auth for user authentication, and *CloudAuthz* for granting cloud authorization.

**Fig. 8.**
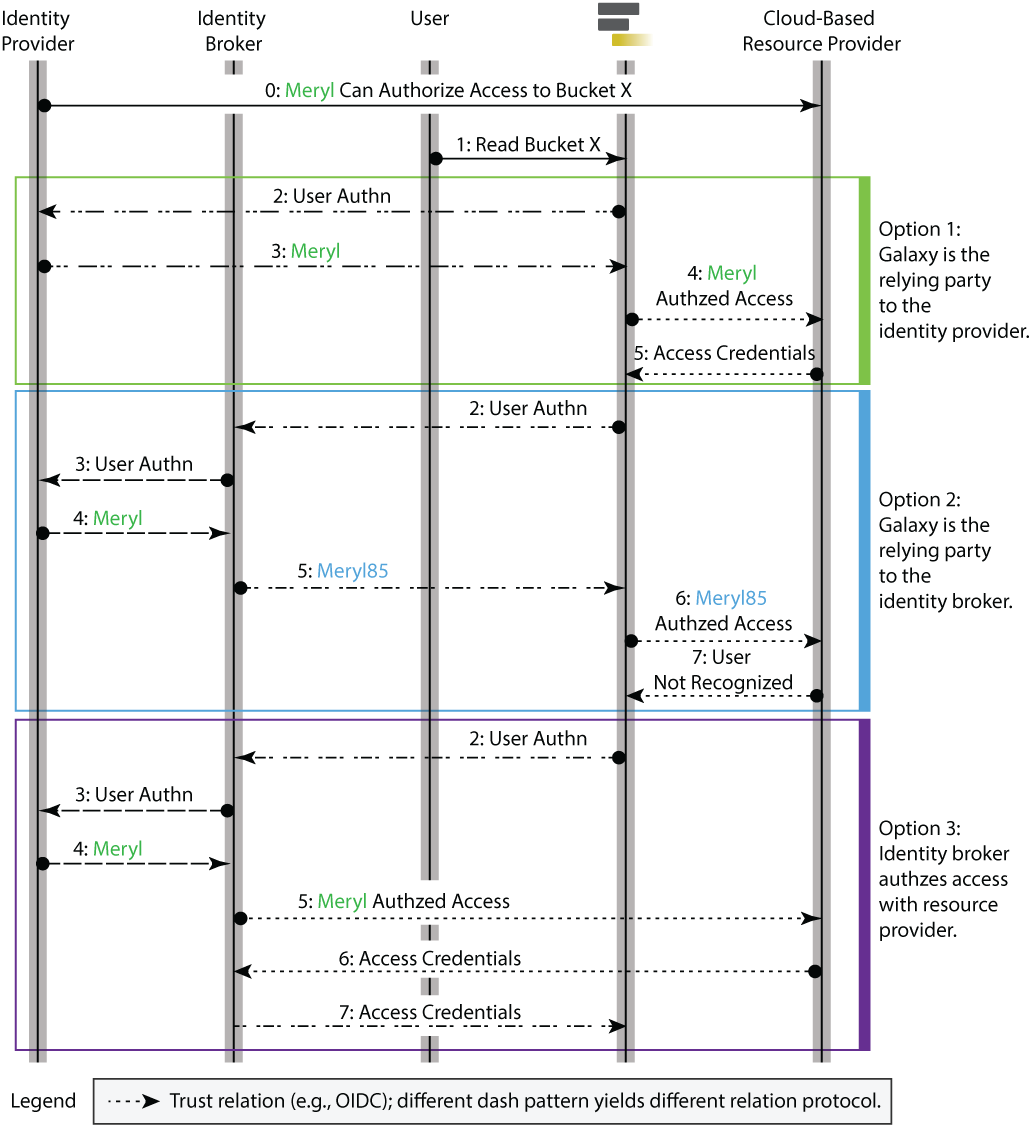
Illustrates three patterns of user authentication and cloud authorization. The *Option 1* is based on *direct* authentication protocol, which we currently implement. The *Option 2* is based on brokered authentication pattern, and since methods implementing this protocol can map an authenticated user to a local identity (see steps 4 and 5 of *Option 2*: the broker emits its own authentication instead of relaying the IdP’s proof), this protocol cannot be used for authorization grant to cloud-based resource providers. In other words, the broker can emit an identity (e.g., “Meryl85”) that is different from the identity expected by the resource provider (e.g., “Meryl”), resulting authorization failure by the resource provider (see step 7 of Option 2). The *Option 3* also follows *brokered* authentication pattern, but since it also provides authorization grant service (Amazon Cognito is such a broker), it can be used as an alternative to *Option 1*.

In the remainder of this section, we discuss the advantages of the proposed method, related security challenges, and compare existing authentication and authorization protocols with our choices.

### A. Countermeasures against eavesdropping attack

The eavesdropping attack is a type of the man-in-the-middle attack where the attacker sniffs and relays communication between parties (e.g., between Galaxy and AWS) and steals sensitive information such as identity tokens and/or access credentials. A common practice that we leverage to effectively thwart eavesdropper revolves around two principles: (1) cryptographically secured communication channel between the parties, and (2) use of OIDC access tokens. Built into the OAuth2.0 protocol, transmitting tokens mandates using Transport Layer Security (TLS) protocol (see sections 10.3 and 10.4 at tools.ietf.org/html/rfc6749). Accordingly, we recommend employing TLS to secure the communication between Galaxy and both identity and resource providers. The OIDC tokens are short-term credentials with least priviledge. Leverage those tokens shortens the time-frame during which an eavesdropper can impersonate a Galaxy user when the TLS connection is exploited and tokens are stolen. The maximum age of tokens and cloud-native credentials is configurable in Galaxy by instance administrators, and we recommend setting it to their minimum values (the default value is 3600 seconds). The `exp` claim of JWTs sets the expiration time of tokens; and since the tokens are cryptographically signed, the value of `exp` claim (among other claims) cannot be changed without invalidating the token. The expired tokens can be refreshed only by trusted parties using their secrets (e.g., *audience* ID and secret) and refresh tokens.

### B. State-of-the-art of Fine-grained Medical Data Access Control in Cloud Computing

As the interest in data-driven healthcare continues to intensify, data security and privacy become imperative, which demands more robust and transparent data governance. Additionally, with the proliferation of biobanks and comparative data analysis methods, data sharing across institutes is becoming essential, which escalates data governance challenges. Siloing of datasets across organizations impedes data usage because accessing each requires a researcher to separately apply for access through a *data access committee*. This ecosystem raises multiple challenges. For example, there has been a great deal of controversy evolving around *consent*. The Global Alliance for Genomics and Health (GA4GH) is fostering “consent codes” to facilitate data sharing, which divides data access conditions into nineteen empirical “categories” and “requirements” for consistent interpretation of data access and consent [15]. However, lack of consensus on the legal and ethical appropriateness of existing strategies hinders their adoption [16].

Another example is determination and automatic enforcement of data usage restrictions and user authorizations. The Data Use Oversight System (duos.broadinstitute.org) is an attempt to define and enforce an ontology of data access restrictions. The GA4GH has launched a pilot study, named “library card” [17], to standardize a role-based authentication and authorization of researchers by augmenting the widely-adopted protocols such as OIDC; it is envisioned to encode “bona fides” of researchers as a set of standardized *claims*. Additionally, GA4GH has defined “registered access”, an empirical data access model that leverages “consent codes” and “library card” for authentication, attestation, and authorization of researchers access to protected data [18],[19]. The different protocols for authentication and authorization are discussed in the following sections.

#### 1) User Authentication Protocols

A principle component to data sharing and their access control is user identification and authentication across institutes and resource/service providers. There has been a number of protocols developed for this purpose, such as Lightweight Directory Access Protocol (LDAP), Security Assertion Markup Language (SAML), OASIS WS-* (Security, Trust, and Federation), OAuth, and OpenID Connect (OIDC). These protocols are used in widely adopted services such as *Shibboleth*, which leverages SAML protocol to enable *single sign-on* across organizations. For instance, using Shibboleth, a university *X* (Shibboleth Identity Provider) with subscription to a publisher *Y* (Shibboleth Service Provider) can enable its students to login to *Y* and access subscription-required articles.

Galaxy supports LDAP, SAML, and now OIDC (see https://galaxyproject.org/authnz/). For the scenarios described in this manuscript, addition of the OIDC protocol was necessary to enable role-based access controls. Namely, the OIDC allows a consistent model for interfacing with major cloud-based resource providers (e.g., AWS, Google Cloud Platform, and Microsoft Azure) when accessing private resources of users by a third party (i.e., Galaxy). Leveraging OIDC-certified libraries (see openid.net/developers/certified/ for their list), any platform can act as an OIDC identity provider; however, to allow users to login to Galaxy using their institutional or social identities, and security challenges of implementing and maintaining an OIDC IdP for developers and Galaxy instance admins (e.g., counterfeits weaknesses covered in tools.ietf.org/html/rfc6819), we are keen to rely on external identities.

There exist two architectural approaches for user authentication using their external identities: *direct* and *brokered* [20], *[21]. In the direct* authentication pattern, a client (e.g, Galaxy instance) directly establishes a *trust* relation (i.e., communicate following a standard protocol such as OIDC) with an IdP, acting as an RP, where the IdP issues identity tokens with `aud` claim being the audience ID of that Galaxy instance. The *direct* authentication is a decentralized pattern, where users are authenticated to different Galaxy instances independently. Using a decentralized pattern, a breached *trust* relation is isolated and cannot affect other *trust* relations. Additionally, admins of Galaxy instances can independently choose IdPs following their institutional policies. However, to interface with multiple IdPs following this pattern, Galaxy needs to implement every IdP-specific *trust* relation.

The *brokered* pattern leverages an authentication broker; an intermediary service of a *single sign-on* architecture that establishes a *trust* relation between multiple IdPs and service providers, and it is *trusted* by both parties independently. In other words, a broker can use different authentication and authorization protocols to communicate with IdP and Galaxy. A broker may decouple parties using an internal user identity, and vouches for the user by issuing its own identity tokens to the clients (see Figure 8) [20]. An advantage of this design is the ability to impersonate a user by masking their login username by the internal identity of the broker to Galaxy. Additionally, an authentication broker can negotiate *trust* between Galaxy and IdPs, which removes the need for direct relation with IdPs. However, the brokered pattern is a centralized approach, where users are authenticated to various Galaxy instances using shared identities. Accordingly, a brokered pattern establishes a single point of failure and a central breach point; a security and liability concern [22]. If compromised, it can jeopardize the security of users on all connected Galaxy instances. If it fails, none of the parties can communicate; however, with the cost of increased design complexity, this problem can be mitigated by significant number of redundant and mirrored brokers. Additionally, some Galaxy instances may not be commissioned to interface brokers or use shared identities due to institutional policies and *consent* concerns.

In spite of the plain core premise of *direct* and *brokered* patterns (as described in [20]), the specification has a myriad of options and variations, which makes it difficult to draw clear boundaries between the available implementations. Some IdPs provide libraries for *direct* user authentication, for instance Google (developers.google.com/identity/protocols/OAuth2) and Microsoft (docs.microsoft.com/en-us/azure/active-directory/develop/reference-v2-libraries). Python Social Auth (github.com/python-social-auth/social-core) implements IdP-specific trust relations for common social identity providers and exposes them via a common interface, which simplifies using the *direct* authentication pattern with multiple IdPs. A decent number of services are available for the *brokered* pattern authentication, spanning from commercial products such as Amazon Cognito (aws.amazon.com/cognito/) and Okta (www.okta.com), to free and open-source services such as Keycloak (www.keycloak.org), CILogon [23]), and Fence (github.com/uc-cdis/fence).

Galaxy needs to authenticate users in a heterogeneous environment of authentication and authorization approaches. An authentication broker can negotiate trust between Galaxy and IdPs (and service providers), which removes the need for direct relation with IdPs. Meanwhile, using libraries such as Python Social Auth, Galaxy can establish a trust relationship with multiple IdPs through a common interface, without the need for an IdP-specific implementation. For the time being, we leverage the *direct* authentication pattern and use the Python Social Auth library to establish a *trust* relation with IdPs.

#### 2) Protocols for User Authorization to Cloud-Based Resources

Cloud-based resource providers commonly implement two methods to authorize access rights to secured resources: *signed URLs* and *cloud-native credentials. Signed URLs* grant a party in possession of the URL particular access to specific resources determined at the URL generation by the resource owner. Signed URLs allow resource owners to grant temporary access to users who are not required to be authenticated by the resource provider. While signed URLs simplify data sharing, it is challenging to audit the access to the data shared using signed URLs. Additionally, the enforcement of a fine-grained access control on a large scale would be challenging since the URLs are generated on a per-resource basis.

*Cloud-native credentials*, as the name implies, are provider-specific secrets to sign programmatic requests to the provider (e.g., API requests). Providers commonly offer long and short-term credentials. A resource owner can obtain long-term secrets from the provider’s portal and use them to authorize a client’s (e.g., a web or native app) access to secured resources. However, such credentials are commonly obtained via a manual intervention that demands a degree of familiarity with the resource provider’s portal. Additionally, the long-term nature of such credentials can encourage scenarios where the tokens are embedded or distributed in applications, which increase the risk of tokens being hijacked.

To address these issues some cloud-based resource providers (e.g., Amazon and Microsoft) implement *security token service* (STS), with the service specifications defined as part of OASIS WS-Trust and WS-Federation protocols [24]. The STS issues short-term security credentials upon a successful assertion of user authentication (see Figure 1) and it is intended to be used by native and web applications as an (OAuth2.0) authorization server. Since such tokens are emitted on-the-fly as per API requests, obtaining them does not require a manual intervention of the resource owner. Additionally, these are short-term credentials with a limited lifetime, thwarting issues described in Section 3.1.

An additional concern is the scope of access for the issued tokens, which should ideally have an option of not being equivalent as the owner’s account. Some resource providers (e.g., Amazon) use federated identities and leverage RBAC model to enable assuming a role using authentications issued for specified clients by determined IdPs (see Figure 4). Leveraging this model, a client can obtain short-term credentials on behalf of users which–critically–are not necessarily part of the resource owner’s cloud subscription account. This is advantageous for sharing data without adding collaborators to a cloud subscription account, for example.

To support this usage models, Galaxy obtains user authorization to protected cloud-based resources leveraging the RBAC model from the resource provider’s STS. Since resource providers expose STS and RBAC differently, we have developed a library called *CloudAuthz*, which provides a common interface to various resource providers. The library currently supports AWS and Azure, with support for the Google Cloud Platform (GCP) under development (see Figure 7). Leveraging CloudAuthz internally allows Galaxy to implement a single interface toward all supported providers.

## IV. Discussion

In this paper, we have described a robust, generalized, and secure approach for an application accessing biomedical data stored on cloud computing platforms on behalf of a user. We use best-practice Web protocols so that user credentials are never requested, transmitted, or stored. We have implemented our approach in the Galaxy platform, which is used across the world for large-scale biomedical analyses. By leveraging role-based access controls and the OIDC protocol, Galaxy users can enable a Galaxy server to access private datasets on a cloud platform using best-practice authentication and authorization approaches. Our approach follows the principle of least privilege and allows a user to revoke Galaxy’s access at any time to one or all datasets. Galaxy’s federated identity and access management features enable users to securely and seamlessly access their protected cloud-hosted (and often sensitive) genomic data and use that data in Galaxy for complex, integrated analyses.

Our work extending Galaxy to use OIDC protocols and RBAC data access to securely access biomedical datasets across cloud platforms is a first step toward developing a user-friendly computational workbench for analysis of distributed biomedical data [7]. Our next step is to implement a *common identity* across Galaxy servers. A common identity paves the path toward a coherent user experience across different Galaxy instances, where users’ data, workflows and analysis histories are unified and accessible from any Galaxy server. A common identity can be implemented by integrating Galaxy with an external user authentication and authorization service such as Keycloak (https://www.keycloak.org/) or Amazon Cognito (https://aws.amazon.com/cognito/).

Another future direction is improving sharing of cloud-based datasets through Galaxy. With our current approach, cloud-based datasets are copied to the Galaxy server, where they can be shared with individual users, by web link, or published to a public list using Galaxy’s collaboration framework. This is problematic because cloud datasets must be replicated to the Galaxy server for sharing, which is time-consuming and can make version and provenance tracking more difficult. We plan to extend Galaxy’s federated identity and access functionality to make it possible for users to share datasets directly via cloud storage by changing the role-based access settings on the cloud storage. Galaxy users could then access shared datasets directly in cloud storage.

Finally, we are adding support for open-source software cloud computing platforms. Globus, Elixir, and Keycloak are being incorporated for user authentication in Galaxy, and Galaxy will soon support OpenStack (https://www.openstack.org/) for RBAC to data on OpenStack. Integrating these platforms will ensure that Galaxy’s federated identity management and data access features can be widely used in academia and other projects that rely on open-source software.

